# First genetic detection and ongoing eDNA monitoring of the golden mussel (*Limnoperna fortunei*) in California

**DOI:** 10.64898/2026.06.18.733028

**Authors:** Sarah Stinson, Aviva Fiske, Emily C. Funk, Emily Kulig, Sarah Brown, Daphne Gille, Andrea Schreier, Leigh Sanders, Raman P. Nagarajan, Bryan Barney, Melinda R. Baerwald

**Affiliations:** California Department of Water Resources, Sacramento, California, USA; Department of Animal Science, University of California, Davis, California, USA; California Department of Fish and Wildlife, California, USA

**Keywords:** invasive species, aquatic invertebrates, genetic monitoring, environmental DNA, San Francisco Estuary

## Abstract

Here, we report the first genetic confirmation of golden mussels (*Limnoperna fortunei*) in North America, and the subsequent development, optimization, and deployment of golden mussel eDNA monitoring procedures. Aquatic species invasions are economically costly, disrupt ecosystem functionality, and impact native aquatic communities. Early detection of new invasive species enables rapid response via implementation of effective eradication or control measures and is key for reducing harmful outcomes. Initial species detection and taxonomic identification can be aided by genetic methods that have high detection sensitivity and accuracy. Genetic methods such as environmental DNA (eDNA) sampling can be used to detect invasive species before they become established in new systems, providing an early alert system to inform resource managers. Golden mussels were first detected in North America in October 2024 near the Port of Stockton in the San Francisco Estuary (SFE). The SFE is particularly vulnerable to invasion due to the access and connectivity provided by the presence of engineering infrastructure and shipping lanes. Collaborative efforts between public agencies and academic institutions are underway to develop a coordinated detection and response plan. Early detection followed by a rapid response is the best defense against prolific invasive species, such as the golden mussel.

## Introduction

Aquatic species invasions negatively impact the global economy, costing billions in US dollars annually (Cuthbert et al. 2021). They also disrupt ecosystem functionality, often altering biodiversity and reducing the abundance of native aquatic communities (Gallardo et al., 2015). Coordinated early detection and rapid response (EDRR) actions can reduce long-term economic and ecosystem consequences by detecting new invasive species and implementing effective eradication or control measures (Reaser et al., 2020). The first steps in the process are initial species detection and taxonomic identification, which can be aided by genetic methods that have high detection sensitivity and accuracy.

Genetic methods such as environmental DNA (eDNA) sampling can be particularly beneficial for the initial detection of invasive species before they become established in new systems. Biological organisms continually shed DNA in multiple potential states, including whole cells (e.g., tissues, waste, and mucus), cellular organelles (e.g., mitochondria), and dissolved or “free” DNA that may also be bound to suspended particles (Mauvisseau et al. 2022). Once shed from an organism, DNA can persist and disperse in the environment in water, soil, or air. eDNA can be collected, then analyzed for presence/absence of a species’ DNA in the environment (Bruce et al. 2021). Sensitive genetic assays can detect minute amounts of eDNA from species that are elusive or present in low abundance at the initial stages of invasion (Dougherty et al. 2016; Mauvisseau et al. 2019; Guo et al. 2024). eDNA surveys often outperform traditional capture-based or visual surveys for invasive species in sensitivity, cost-per-unit effort, and speed, which enables surveillance of large areas and potentially inaccessible habitat with a representative environmental sample (Fediajevaite et al. 2021; Rishan et al. 2023). eDNA sampling is also relatively straightforward and can be performed by community scientists to enhance invasive species detection and monitoring (Meyer et al. 2021). Alternatively, if a biological organism is collected, genetic methods (e.g., mitochondrial DNA analysis) can rapidly and reliably identify an emerging invasive species, even when morphological differentiation is challenging due to cryptic speciation (Darling and Blum 2007; Li et al. 2024).

The golden mussel (*Limnoperna fortunei*) is a freshwater bivalve native to China that is known to rapidly invade susceptible ecosystems (Xia et al. 2017; Boltovskoy et al. 2025). Once established, this invader acts as an ecosystem engineer, physically altering habitat structure and function via direct biofouling, altering oxygen availability and promoting toxic cyanobacterial blooms (Darrigran and Damborena 2011; Haubrock et al. 2022). Other ecological impacts include food web disruption, rapid depletion of phytoplankton and zooplankton species, changes in the composition of benthic invertebrate species, and promotion of certain fish parasites (Boltovsky and Correa 2015; Cataldo et al. 2012). Economic impacts can result from extensive biofouling of water supply infrastructure, obstructing flow, impacting water supply sources for drinking water treatment plants, fire protection systems and power plants (Carranza et al. 2023), impaired water conveyance capability, structural damage, and water quality contamination (Yang et al. 2024, Boltovskoy et al. 2024, Ostrensky et al. 2024). The species has spread to several countries in Asia (e.g., Japan, Korea, Thailand) and South America (e.g., Argentina and Brazil) (Boltovskoy et al. 2025), causing extensive damage both to water infrastructure and ecosystem function (Haubrock et al. 2022; Heringer et al. 2021). As of 2022, the total economic cost of *Limnoperna* global invasions was estimated at $140.5 million USD (Haubrock et al. 2022), representing a significant portion of all freshwater bivalve invasions.

Even twenty-five years after the initial invasion, the annual economic loss in Brazil is still approximately $9.97 million USD (Ostrensky et al. 2025, Heringer et al. 2021). Golden mussels were recently detected in North America near the Port of Stockton in the San Francisco Estuary (SFE) during a routine water quality station check. The SFE is particularly vulnerable to invasion due in part to the access and connectivity provided by the presence of engineering infrastructure (i.e., the State Water Project) and heavily trafficked shipping ports (Galil et al. 2008; Zhan et al. 2015). The SWP is a multi-purpose water storage and delivery system that delivers clean water to 27 million Californians and 750,000 acres of farmland, while supporting flood control efforts (Seckler, 1971).

### A brief timeline of golden mussel invasions

The invasion timeline for golden mussels prior to entering North America has been well documented (Boltovskoy et al. 2025), and notable advances are included in Figure 1a. The timeline of the North American invasion is shown in Figure 1b. On October 17, 2024, California Department of Water Resources (DWR) staff observed mussels during a routine field survey in Stockton, CA. Samples were subsequently sent to the California Department of Food and Agriculture (CDFA) Plant Pest Diagnostics Center, the Smithsonian Environmental Research Center, and the UC Davis Genomic Variation Laboratory (Davis, CA). The Smithsonian Environmental Research Center (SERC) shared an initial assessment on October 21, 2024, that the mussels were visually consistent with a *Limnoperna* species, such as *L. fortunei*. On October 23, 2024, the UC Davis Genomic Variation Laboratory genetically confirmed that the unidentified mussel was indeed the golden mussel, *L. fortunei*. CDFA also performed a genetic confirmation which was reported on the same day. The combined results from DNA sequencing and morphological examination confirmed the first reported observation of golden mussels in North America. On October 25, 2024, California State Parks (State Parks) staff detected unknown mussels on an artificial substrate in O’Neill Forebay, downstream of the Delta, which were subsequently confirmed as golden mussels. Shortly thereafter, golden mussels were detected in the State Water Project (SWP) (visually observed on October 24, 2024, genetically confirmed on October 28, 2024, then detected from eDNA samples on December 10, 2024).

**Figure 1.**
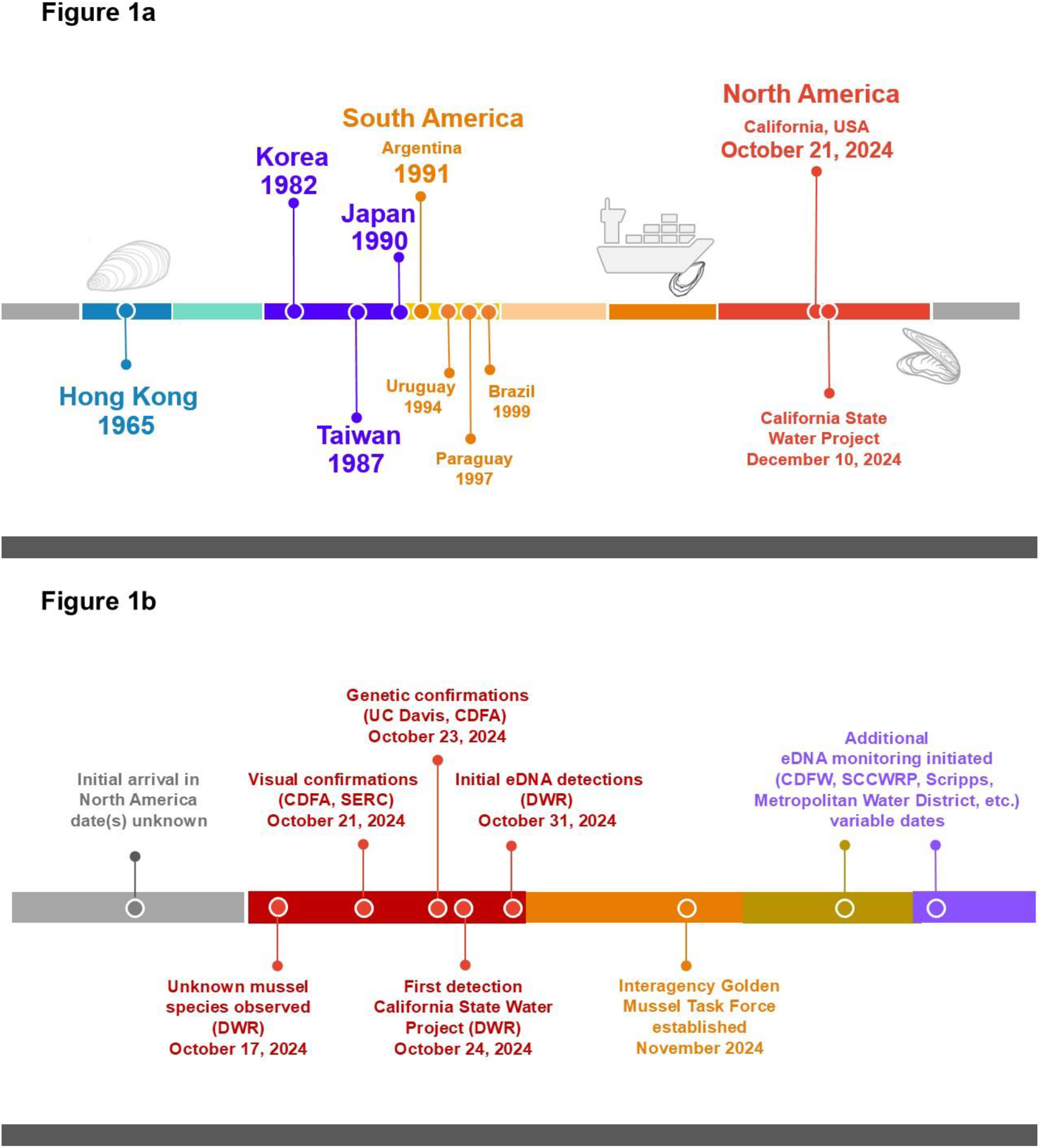
The global invasion timeline for golden mussels (a) as previously reported in the literature, and (b) the invasion timeline for golden mussels entering North America, described herein.

To monitor and assess the spread of golden mussels in California and beyond, an interagency Golden Mussel Task Force (GMTF) is developing a coordinated detection and response plan, following established EDRR processes. Herein, we describe the use of genetic methods to confirm the initial visual identification of the species after it was first observed in the SFE. We also report the integration of eDNA techniques into existing monitoring efforts to assess both spatial (i.e., *where* are golden mussels found?) and temporal (i.e., *when* do golden mussels first appear in an area?) characteristics of the SFE invasion.

## Methods

### Genetic Species Identification

Mussels were collected by California Department of Water Resources (DWR) staff on October 21st, 2024, from the Port of Stockton (Rough and Ready Island, Stockton, CA) and delivered to the Genomic Variation Lab at UC Davis (Davis, CA) for genetic identification. We extracted 6 mussel specimens using the Qiagen DNeasy Blood & Tissue Kit (QIAGEN Redwood City, CA) following the manufacturer’s protocol. We then PCR amplified the cytochrome c oxidase subunit 1 (CO1) gene using degenerate primers, mlCOIintF_adapt and jgHCO2198_adapt (Geller et al. 2013, Leray et al. 2013, Folmer et al. 1994) and visualized the product using gel electrophoresis. We successfully amplified the CO1 gene from 4 of the 6 mussel specimens, and we sent the PCR product from those 4 specimens to Quintara Biosciences (Cambridge, MA) for Sanger sequencing.

### eDNA Assay Specificity and Sensitivity

To ensure species specificity and rule out potential cross-amplification, we tested the Pie et al. (2017) golden mussel qPCR assay on a panel of 10 endemic and invasive bivalve species known to be present in the California Bay-Delta (Table 1). Samples were provided by the California Academy of Sciences Invertebrate Zoology Collections and UC Davis. Where samples from multiple individuals of a species were available, we pooled the extracted DNA prior to qPCR to maximize our chance of detection and minimize risk of false negative results. Three technical replicates for each species were included in the plate, along with no template controls (NTCs) and tissue positive controls. For subsequent tests, synthetic DNA (gBlock™) was used for positive controls. We used qPCR primers and probe reported in Pie et al. 2017 (Table 2), however cycling conditions and primer/probe concentrations were modified significantly to improve assay sensitivity (see below).

**Table 1.** 10 endemic and invasive bivalve species known to be present in the California Bay-Delta. Extracted DNA from these species was used to test the specificity of the Pie et al. 2017 qPCR assay. Samples were provided by the California Academy of Sciences Invertebrate Zoology Collections and UC Davis.

Endemic bivalves of the San Francisco Bay Delta (CA, USA)
| Scientific name | Common name(s) |
| --- | --- |
| *Andodonta californiensis* | California floater |
| *Gonidea angulata* | western ridged mussel |
| *Margaritifera falcata* | western pearlshell |
| *Mytilus californianus* | California mussel |
| *Mytilus galloprovincialis/trossulus*<br>*complex* | Pacific Blue mussel/Foolish mussel/Mediterranean mussel |

Invasive bivalves of interest
| Scientific name | Common name(s) |
| --- | --- |
| *Dreissena polymorpha* | Zebra mussel |
| *Dreissena bugensis* | Quagga Mussel |
| *Musculista senhousia (San Diego)* | Asian date mussel/Asian mussel/bag mussel |
| *Corbicula fluminea* | Asian Freshwater Clam/Golden clam/freshwater golden clam |
| *Potamocorbula amurensis* | overbite clam/Asian clam/Amur River clam |

**Table 2.**
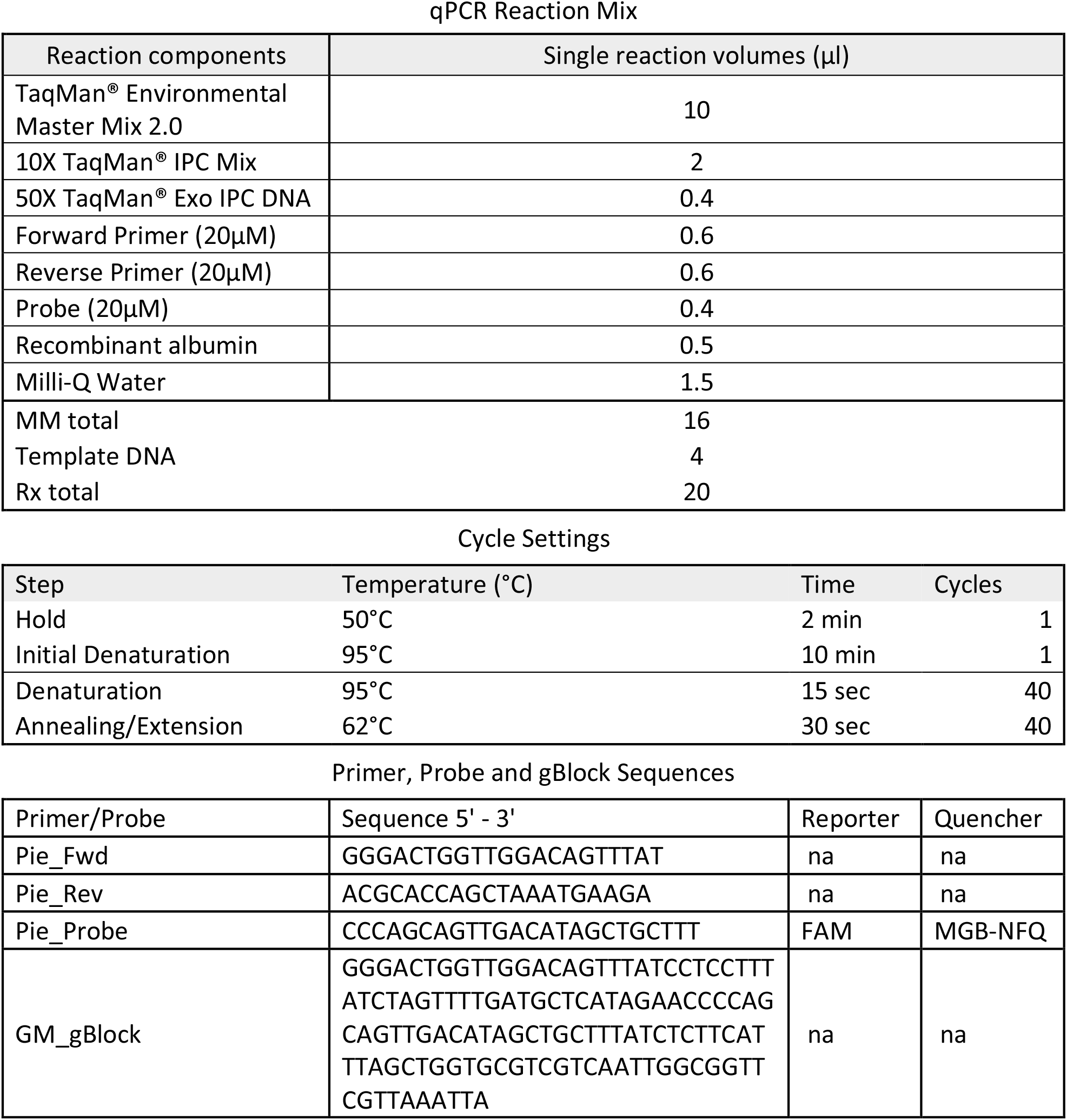
Concentrations and volumes of reagents used in the qPCR reaction, qPCR cycle settings and sequences (5’-3’) of the forward and reverse primers, probe and gBlock are listed.

Authors at the California Department of Fish and Wildlife (CDFW) Genetic Research Laboratory and the DWR Genetic Monitoring Laboratory performed a series of experiments to optimize the sensitivity of the qPCR assay, as well as laboratory proficiency tests to ensure comparable results. We designed synthetic double stranded DNA (gBlock™) from the mitochondrial sequence of the CO1 gene in the golden mussel reported in NCBI GenBank accession DQ264395.1 to create our standard curves and quantify the DNA molecules of interest (Integrated DNA Technologies, Coralville, ID). Reaction chemistry and cycle settings for the qPCR assay are shown in Table 2. The solution was prepared in a 20 µl volume including 10 µl of TaqMan® Environmental Master Mix 2.0 (Applied Biosystems, Waltham, MA), 2 µl of TaqMan® exogenous 10X mix, 0.4 µl TaqMan® exogenous internal positive control DNA (IPC, Applied Biosystems, Waltham, MA), 0.5 µl of molecular grade Recombinant Albumin (New England Biolabs, Ipswich, MA), 4 µl of DNA template, 600 nM of each primer, and 400 nM of probe, and 1.5 µl of molecular-grade water (Applied Biosystems, Waltham, MA). The reaction was performed at CDFW and DWR using QuantStudio 5 or 6 (Applied Biosystems, Waltham, MA), respectively. Reactions were performed using standard settings with a hold at 50°C for 2 minutes then 95°C for 10 minutes, followed by 40 cycles of denaturation at 95°C for 15 seconds, and annealing/extension at 62°C for 30 seconds. No template controls (NTCs) and gBlock™ standard curves were included on all qPCR plates. Samples were required to have Cq values below the threshold (before 40 cycles) from at least 2 of 3 technical replicates to be considered positive. Any samples where 1 of 3 replicates amplified were re-run for confirmation. We used the discrete method described in Klymus et al. 2020 to determine the limit of detection, limit of quantification, efficiency and R2 values, using the R environment (R Core Team 2026, version 4.1.3).

### Ongoing eDNA monitoring

DWR tow net samples are collected routinely on a year-round basis from water quality stations throughout the Bay-Delta. For this study, all samples were collected either as 1 L water grab samples or using tow nets (Sea-Gear Corporation, Florida USA). To obtain tow net samples, a 45-meter horizontal surface net tow was performed at each site. Samples were rinsed into, then collected from, the cod end of the net (64 μm mesh) and immediately placed on ice. Samples were preserved with > 50% by volume of molecular grade 200 proof ethanol (ThermoFisher Scientific, Waltham, MA). Nets and equipment were decontaminated with 10% bleach (submerged for > 5 minutes), then rinsed thoroughly with deionized water. Water grab samples were collected by submerging a sterile 1 L container (ThermoFisher Scientific, Waltham, MA) in site water, then storing on ice until filtration using a GeoTech peristaltic pump (GeoTech Environmental Equipment, Inc. Sacramento, CA) and 47 mm GF filters (ThermoFisher Scientific, Waltham, MA). Field blanks were collected (immediately prior to sampling, under field conditions) by filtering deionized water through the tow net, collecting 25 mL from the cod end, and preserving it in > 50% by volume of molecular grade ethanol in sterile 50 mL Falcon conical centrifuge tubes (Corning Inc., Sunnyvale, CA). Ice chest blanks consisted of sterile 50 mL tubes containing 25 mL deionized water, placed directly into the ice chest used to store samples and field blanks. Field blanks and ice chest blanks were collected for every sampling event. Samples were placed on ice and transported to the lab for processing. All samples were frozen or processed within 12 hours of submission.

We extracted eDNA from tow net samples following Schabaker et al. 2020. All eDNA sample processing was performed in the DWR Genetic Monitoring lab (West Sacramento, CA), in our dedicated eDNA laboratory space (all procedures were completed in PCR hoods with UV lights, the space contains no amplified DNA products, and decontamination procedures for surfaces are performed using either bleach or RNase AWAY). We treated all extracted DNA with a OneStep PCR Inhibitor Removal Kit (Zymo Research, Irvine, CA) to purify DNA following the manufacturer’s protocol. Extraction blanks were included in all sample processing events. DNA quantification was performed on a Qubit 4.0 fluorimeter (ThermoFisher Scientific, Waltham, MA). Quantitative PCR was performed on a QuantStudio 6 (ThermoFisher Scientific, Waltham, MA). Field blanks, extractions blanks, NTCs, IPCs and gBlock™ standard curves were included on all qPCR plates.

### Mapping the spread

The DWR Genetic Monitoring Laboratory and the CDFW Genetics Research Laboratory are conducting routine qPCR analysis from eDNA samples collected throughout the SFE to map the spread of the golden mussel invasion. Golden mussel sampling locations and detections from December 11^th^, 2023 (archived samples) to December 10^th^, 2025 were mapped using the package ‘leaflet’ (Cheng et al. 2019) in the R environment (version 4.4.3).

## Results

### Genetic Species Identification

Sanger sequencing with the forward primer produced clear and reliable sequence data, but the reverse primer did not; therefore, we only used the results from sequencing in the forward direction for genetic identification. We blasted our sequencing results against the NCBI nucleotide database, and on October 23, 2024, we identified all specimens to be golden mussel (*Limnoperna fortunei*) with 100% identity (raw data is available in the Supplemental data).

### eDNA Assay Specificity and Sensitivity

Specificity testing of the Pie et al. qPCR assay resulted in no amplification of other bivalve species known to be present in the SFE (listed in Table 1), and positive and negative controls performed as expected.

Authors from the DWR Genetic Monitoring and CDFW Genetics Laboratories performed a series of experiments to optimize the sensitivity of the qPCR assay, as well as blind interlab sample exchanges to ensure comparable results. Through this process, we determined that, in contrast to the original assay publication, different reaction concentration and cycle settings (see Methods for details) were necessary to achieve satisfactory efficiency and LOD/LOQ values. The assay standard curve, LOD, LOQ, efficiency and R^2^ for the optimized Pie et al. assay are reported in Supplemental Figure 1. After optimization, the LOQ was 7.0 copies per qPCR reaction, and the LOD was 4.17 copies per qPCR reaction.

### Ongoing eDNA monitoring

A total of 400 eDNA samples were tested, with collection dates ranging from December 11^th^, 2023 (archived samples) to December 10^th^, 2025 (Figure 2). Of that total, 49 samples were positive for golden mussel DNA and 351 were negative. The first positive eDNA detections in North America were obtained from samples collected from sites routinely monitored by DWR in the South Delta on October 31^st^, 2024 (Byron, Light Tower 18, Victoria island, Turner cut, Bacon island, and Buckley cove).

**Figure 2.**
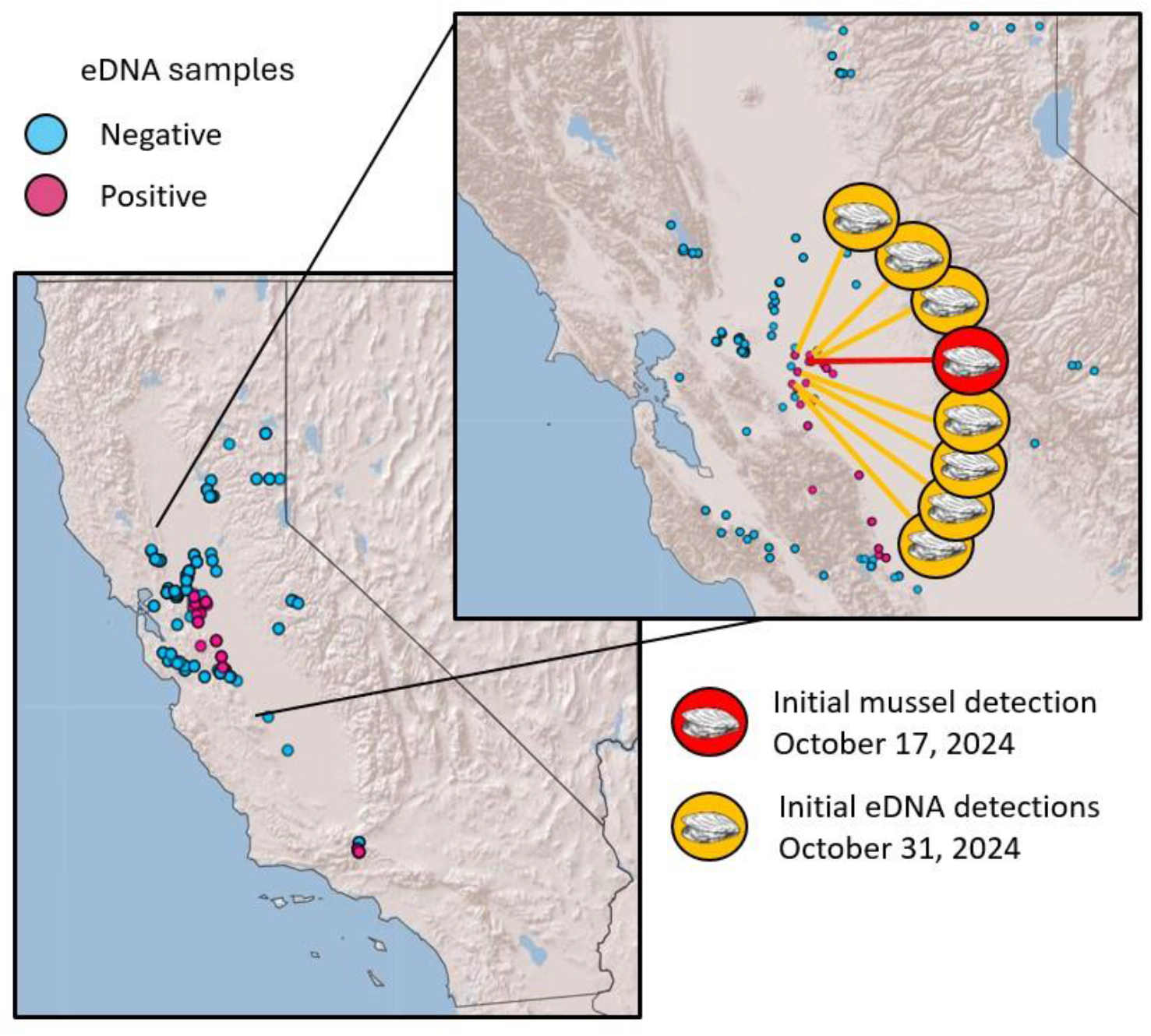
Map of eDNA golden mussel sampling locations (California, USA) with collection dates from December 11^th^, 2023 (archived samples) to December 10^th^, 2025. Blue circles indicate golden mussels were not detected via eDNA (N = 351), and pink circles indicate that golden mussels were detected via eDNA (N = 49). The site of the initial observation and detection of mussels is indicated by a red circle/mussel icon. The sites of the first eDNA detections are indicated by yellow circle/mussel icons.

The Golden Mussel Task Force (GMTF) is an interagency (California Department of Fish and Wildlife, California State Parks, California Department of Water Resources, California State Lands Commission, California Department of Food and Agriculture, California State Water Resources Control Board, United States Bureau of Reclamation, United States Fish and Wildlife Service) effort to develop a coordinated detection and response plan, using eDNA detections as a primary monitoring tool to focus response efforts and maximize available resources. These collaborative efforts are part of a larger movement to transition eDNA technology from Research to Routine Application (Lee et al. 2024). The GMTF has developed a Response Plan to provide recommendations to policy makers, managers, and the general public on how to respond to the recent detections of golden mussel in a quick and effective manner through a common framework across agencies and jurisdictions. It is the goal of the State of California to prevent further introduction and spread of golden mussel within the state, contain mussels within currently infested waters, and eradicate mussels from infested waters where feasible.

The GMTF has identified eight objectives that provide a targeted response to ensuring the prevention, detection, and containment of golden mussel in California. These objectives cover media communication, education and outreach, containment at infested waters, prevention at uninfested waters, monitoring for the presence of golden mussel (including early detection of adults and veligers, and eDNA monitoring), partner engagement, population suppression, science and capacity building. To support this effort, the DWR Genetic Monitoring Laboratory and the CDFW Genetics Research Laboratory are conducting routine qPCR analysis from eDNA samples collected throughout the SFE to map the spread of the golden mussel invasion (Figure 2). This map is routinely updated and available to resource managers, to track the spread of the invasion.

## Discussion

Our results represent the first genetic identification and eDNA detections of the golden mussel in North America. Results from the collaborative monitoring efforts conducted by DWR and CDFW are available to resource managers to monitor the spread of the invasion. These findings highlight the potential power of eDNA sampling as an early warning system and monitoring tool for tracking invasive species to inform adaptive management strategies (Sepulveda et al. 2023). Considered along with other early detections of golden mussel eDNA reported globally (Xia et al. 2017), our results support the implementation of routine eDNA monitoring for golden mussels, as well as other potential SFE invasives, in California and beyond.

As with any monitoring tool, there are factors affecting detection probability which should be considered when planning an eDNA-based monitoring strategy. The abundance of golden mussel eDNA may increase during the reproductive season, when gametes are expressed and veligers are present. The length of the golden mussel reproductive season is mediated by water temperature (Boltovskoy et al. 2025), and therefore, at optimal temperatures, spawning activity- and thus detection probability-may increase. Concurrently, temperature and other physicochemical and hydrological conditions including flow, pH, dissolved O_2_, and salinity can influence the state of preservation of eDNA molecules, which in turn influences detection probabilities (Barnes & Turner 2016). These factors may also influence the presence of golden mussels and their reproductive output. Species-species interactions may also affect detection

(Thompson & Gallager 2012). For example, veligers are vulnerable to toxic microcystins and may be absent from locations where microcystin-producing cyanobacterial blooms have recently occurred (Boltovskoy et al. 2013). For eDNA studies, care should be taken to check that qPCR primers are appropriate for detecting local populations, which may carry population-specific genetic haplotypes that reduce qPCR efficiency (Adams et al. 2019; Pinfield et al. 2019).

Golden mussels are now present in California and have begun to disperse through waterways including the SWP. Based on the timeline of invasion and rate of spread observed in the South American invasion, they will be extremely difficult, if not impossible, to eradicate completely. Monitoring will continue to track their presence in the SFE and evaluate the success of control measures. When eDNA monitoring is routinely used as part of an early warning system, resource managers gain valuable time to mount a response. Whether additional time would have prevented their advance into the SWP is unknown, as the connectivity and accessibility of the system make it extremely vulnerable to invasion. However, a proactive monitoring approach may provide additional warning for other potential invaders. We propose that developing genetic assays (i.e., either targeted qPCR or metabarcoding) capable of detecting a “most wanted” list of potential invasives could be invaluable for rapid detection. The added value gained from reanalysis of archived samples and/or leveraging samples being collected for other programs could provide valuable insight on the timing and spread of invasives. Interagency collaborative efforts, as illustrated by this study, have the potential to maximize all available resources and amplify the capacity for rapid response to invasive species.

## Statements and Declarations

### Funding Statement

The authors declare that no funds, grants, or other support were received during the preparation of this manuscript. The work was internally funded by the authors’ respective agencies and no external funds were received for the work.

### Competing Interests

The authors have no relevant financial or non-financial interests to disclose.

### Author Contributions

Study conception and design were performed by all authors. Material preparation, data collection and analysis were performed by Sarah Stinson, Aviva Fiske, Emily C. Funk, Leigh Sanders, Emily Kulig, and Bryan Barney. The first draft of the manuscript was written by Sarah Stinson, Daphne Gille, Melinda Baerwald, Emily Funk and Emily Kulig, and all authors commented on subsequent versions of the manuscript. All authors read and approved the final manuscript.

## Acknowledgements

Sample collections: California Academy of Sciences Invertebrate Zoology Collections staff, California Department of Water Resources Environmental Monitoring Program (Erin Pomidor, Jay Aldrich, Scott Waller), University of California at Davis (Todgham lab, Alexis Simon, Coop lab); Morphological and genetic identifications: California Department of Food and Agriculture (Kevin Williams, Doris Yu, Peter Kerr), Smithsonian Environmental Research Center (Shawn McBride, Andrew Chang, Greg Ruiz, Greg Ziegler, Katrina Lohan), California Department of Parks and Recreation (Lee Sencenbaugh); Consultation & discussion: California Department of Water Resources Operations and Maintenance (Brianne Sakata, Tanya Veldhuizen), California Department of Water Resources Division of Science and Engineering (Alison Collins, Karen Gehrts, Bryan Nguyen), Interagency Golden Mussel Task Force eDNA sub-team (Thomas Jabusch, Susanna Theroux, George Di Giovanni, Jaque Keele, Anthea Lee, Yale Passamaneck, Adam Sepulveda). This work was supported by funding from the California Department of Water Resources.

## Notes

### Competing Interest Statement

The authors have declared no competing interest.

