## Supplemental-Figure-1 for "First genetic detection and ongoing eDNA monitoring of the golden mussel (*Limnoperna fortunei*) in California"

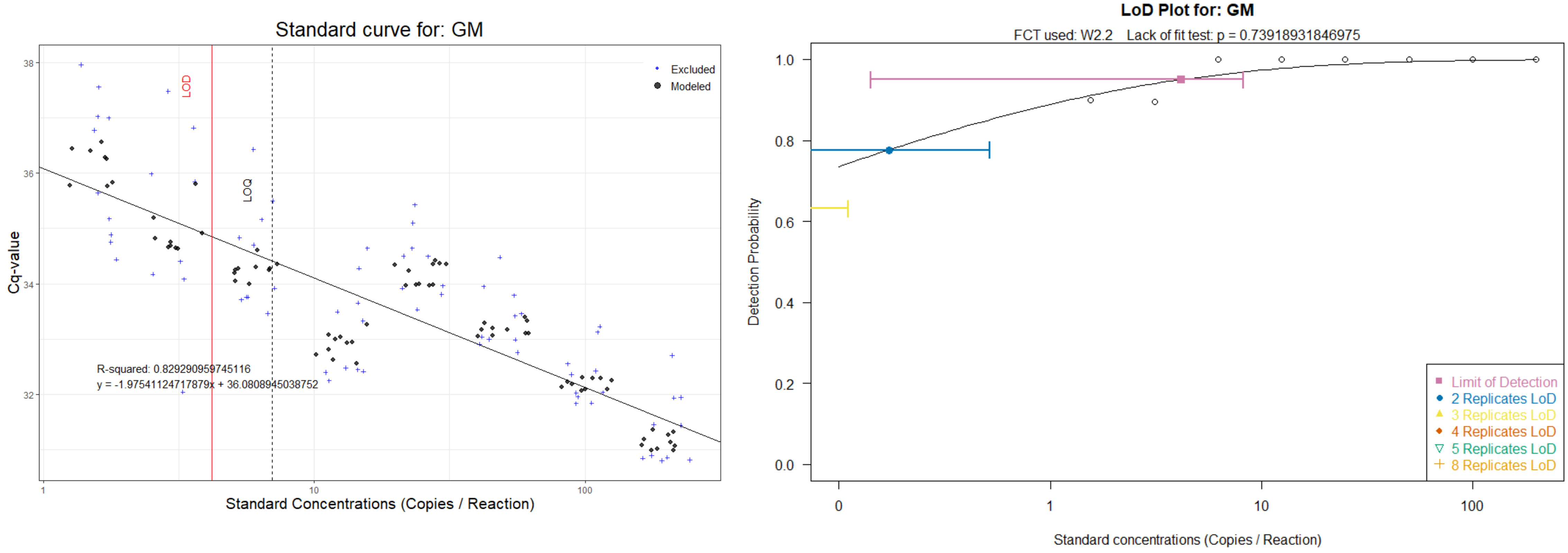


| R.squared | Slope | Intercept | Low.95 | LOD | LOQ |
|---|---|---|---|---|---|
| 0.829291 | -1.9754 | 36.0808945 | 6.25 | 4.17105 | 7 |

Supplemental Figure 1: The assay standard curve, LOD, LOQ, Efficiency and $R^2$ for the optimized Pie et al. assay were determined using R scripts developed by Klymus et al. 2020 in the R environment (version 4.1.3).
